# Sex differences in interrelationships between arterial stiffness, carotid intima-media thickness, white matter hyperintensities, depression and cognition: A UK Biobank study

**DOI:** 10.1101/755280

**Authors:** Ruby S. M. Tsang, John E. Gallacher, Sarah Bauermeister

## Abstract

**Objective:** To explore sex differences in the associations between arterial stiffness index, carotid intima-media thickness, white matter hyperintensities, depression and cognition.

**Methods:** UK Biobank is a population-based cohort study of 502,664 healthy community dwelling adults aged 37-73 years. A select number of participants were recalled to participate in an online reassessment and imaging study, both of which included repeat cognitive assessments. A total of 7,394 volunteers aged 45-73 years (55% female) participated in the imaging visit and completed the self-report mental health questionnaire in the online follow-up were included in the analyses reported here. The main outcome measure of depression was measured using the PHQ-9 and cognition was assessed through measures of reaction time, verbal-numeric reasoning and visual memory. Pulse wave velocity (PWV) was assessed non-invasively using finger photoplethysmography, carotid intima-media thickness (CIMT) with automated ultrasound, and white matter hyperintensity volume with combined T1 and T2-weighted fluid-attenuated inversion recovery (FLAIR) magnetic resonance imaging (MRI).

**Results:** Cross-sectionally, greater arterial stiffness was associated with greater depression in men but with better cognition in women. When white matter hyperintensities burden was added to the model, it mediated the relationships of carotid intima-media thickness with depression and cognition only in men.

**Conclusions:** We report sex differences in brain microvascular changes, depression and cognition in ageing, and suggest that they may be partly explained by sex-specific effects of vascular ageing.

**Summary boxes:** **Section 1: What is already known on this topic**

- Arterial stiffness and carotid intima-media thickness are two non-invasive vascular ageing markers that have been shown to be associated with depression, cognitive impairment and dementia.
- Some studies report sex differences in arterial stiffness and carotid intima-media thickness.
- There is, however, a paucity of research on sex differences in the associations between these vascular ageing markers, white matter hyperintensities, depression and cognition.

**Section 2: What this study adds**

- Cross-sectionally, greater arterial stiffness was associated with greater depression in men but with better cognition in women. When white matter hyperintensities burden was added to the model, it mediated the relationships of carotid intima-media thickness with depression and cognition only in men.
- Our findings add to the existing evidence base of sex differences in brain microvascular changes, depression and cognition in ageing, and suggest that they may be partly explained by sex-specific effects of vascular ageing.

## Introduction

The contribution of vascular factors to cognitive impairment and dementia has long been recognised^1 2^. Arterial stiffness and carotid intima-media thickness (CIMT) are non-invasive markers of vascular ageing, the former reflects the gradual fragmentation and loss of elastin and accumulation of collagen in the large arteries during ageing, and the latter is a marker of subclinical atherosclerosis. While these markers represent separate processes in vascular ageing, both are independent predictors of vascular events including myocardial infarction and stroke^3 4^, with growing evidence that they may be related to cognitive impairment and dementia^5–8^. Furthermore, such vascular ageing changes may also underlie depression^9–11^, which is a known major risk factor for cognitive impairment and dementia^12 13^. These vascular ageing changes are thought to lead to depression and cognitive impairment through cerebral hypoperfusion, which in turn results in microvascular ischaemia and the development of white matter hyperintensities (WMHs)^14 15^.

WMHs are increased signal intensities on T2-weighted fluid-attenuated inversion recovery (FLAIR) magnetic resonance imaging (MRI). Both the prevalence and severity of WMHs increase with age, and over 90% of older adults show some degree of white matter lesions^16^. The severity and localisation of these WMHs appear to be dependent on age and sex; with women having more of both subcortical and periventricular WMHs, particularly in the frontal region^16^. More recent research suggests that lesion localisation may have topographically specific effects on affective symptoms and cognition. The severity of periventricular WMHs tend to be associated with psychomotor speed^17^ as well as attentional deficits in individuals with late-life depression^18^, whereas deep WMHs are associated with depressed affect^19^ and motivation^20^. A recent study also showed that WMHs close to the frontal horns are mainly associated with executive function performance, parieto-temporal WMHs close to the posterior horns with memory performance, and WMHs in the upper deep white matter with motor speed performance^21^. While the exact pathogenesis of WMHs is poorly understood, histopathological studies show the periventricular and deep WMHs are differentially influenced by vascular factors, with the former more influenced by haemodynamic mechanisms and the latter by small vessel ischaemia^11^.

Current research suggests that there are also some sex differences in vascular ageing^22^, with studies reporting greater arterial stiffness in women than in men^23–25^, but greater CIMT in men than in women^26 27^. However, few studies investigating vascular contributions to depression or cognition in mid- to late-life have systematically explored sex differences^28 29^, raising the possibility that important sex-specific associations have been overlooked. The aim of this study is to explore sex differences in the associations between arterial stiffness, CIMT, WMHs, depression, and cognition.

## Methods

### Participants

UK Biobank is a large population-based cohort study. Over 500,000 participants, aged 37-73 years, were recruited at 22 assessment centres across the UK between 2006 and 2010. The assessment comprised extensive self-report questionnaires, physical, cognitive and functional measures, and collection of biological samples. A more detailed description of the methodology used and data collected in UK Biobank has been reported elsewhere^30^.

The present analysis is restricted to a subsample of UK Biobank participants who participated in the imaging visit and also completed the self-report mental health questionnaire in the online follow-up. Participants with missing data on any of the variables of interest were excluded from the analysis. Further, those who self-reported neurological system cancers, neurological problems that may affect cognition or mood, or psychotic/manic disorders were also excluded. Those with stroke or dementia, however, were not excluded to avoid introducing a collider bias. A list of self-reported illnesses used as exclusion criteria are provided in the **Supplementary Information**.

### Variables

#### Arterial stiffness

Pulse wave velocity (PWV) was assessed non-invasively using finger photoplethysmography (PulseTrace PCA2, CareFusion, San Diego, CA) over 10-15 seconds. An arterial stiffness index (ASI, in m/s) was computed by dividing standing height of the participant by the time between the systolic and diastolic wave peaks. An earlier study has shown that the stiffness index estimated from the contour of the digital volume pulse is highly correlated with carotid-femoral PWV (*r* = 0.65)^31^, which is currently the gold standard measure of arterial stiffness and a strong independent predictor of all-cause and cardiovascular mortality. A higher ASI reflects stiffer large arteries. ASI values beyond 3 standard deviations from the mean were considered outliers and excluded, which resulted in a loss of 0.058% of data points.

#### Carotid intima-media thickness

Automated CIMT ultrasound measurements were performed using CardioHealth Station (Panasonic Biomedical Sales Europe B.V., Leicestershire, UK), with a 5-13MHz linear array transducer. Firstly, a two-dimensional (2D) scan of both carotids was performed along the short axis (transverse plane) from below the carotid bifurcation to below the jaw, with the right carotid scanned first. The scan is stored as a cine loop. Then this was repeated in the long axis (longitudinal plane). Following the 2D scan, CIMT measurements were performed two pre-defined angles for each carotid: right 150°, right 120°, left 210°, and left 240°. CIMT was computed as the mean of the four mean IMT measurements and then natural log-transformed. Participants with one or more invalid scans were excluded. Increased common CIMT reflects very early atherosclerotic changes and is associated with an unfavourable cardiovascular risk profile. Further information on the imaging protocol and quality assurance procedures is described elsewhere^32^.

#### White matter hyperintensities volume

Our study made use of imaging-derived phenotypes generated by an image-processing pipeline developed and run on behalf of UK Biobank^33^. Briefly, WMHs were automatically segmented from the combined T1 and T2-weighted fluid-attenuated inversion recovery images using the Brain Intensity Abnormality Classification Algorithm tool^34^. The WMHs volumes (in mm^3^) were expressed as the percentage of total intracranial volume (i.e. the sum of grey matter, white matter and ventricular cerebrospinal fluid volumes) and then natural log-transformed.

#### Reaction time

Reaction time (RT) was measured using a computerised ‘Snap’ card game. Participants were shown two cards with symbols on the touchscreen, and asked to press a button as quickly as possible when the symbols on the cards shown match each other. This task involved 12 trials, with the first five trials regarded as practice trials. The score on this task was the mean response time (in milliseconds) across trials that displayed matching cards. An earlier study reported that these trials showed high internal consistency (Cronbach’s *α* = 0.85)^35^. Unusually fast (<150ms) or unusually slow (>2000ms) responses were excluded, which resulted in the loss of 0.042% of data points across the four matching trials. We computed an intraindividual mean RT for all participants who have valid data from three or more trials, which was then natural log-transformed.

#### Visual memory

Visual memory was assessed using a pairs matching game. A set of cards with symbols were randomly displayed on the screen before they were turned over. Participants were asked to memorise the positions of the cards, and match them from memory while making as few errors as possible. The first round involved three matching pairs shown for three seconds, and the second round involved six matching pairs shown for five seconds. The score on this task was the total number of incorrect matches across the two rounds, which was natural log-transformed. Further details of the cognitive tasks used in the UK Biobank have already been reported elsewhere^36^.

#### Verbal-numerical reasoning

Verbal-numerical reasoning ability were assessed using a task of 13 multiple choice logic and reasoning items with a two-minute time limit. The score on this task is the number of correct items. The Cronbach’s alpha for these items has been previously reported as 0.62^35^.

#### Depression

Depression was assessed using the self-report nine-item Patient Health Questionnaire (PHQ-9)^37^. Participants rated whether they had been bothered by each of nine presented depressive symptoms in the previous two weeks on a scale from zero to three, corresponding to the categories of “not at all”, “several days”, “more than half the days”, and “nearly every day”.

#### Education

Participants self-reported the qualifications they hold among the options of “college or university degree”, “A-levels/AS levels or equivalent”, “O-levels/GCSEs or equivalent”, “CSEs or equivalent”, “NVQ or HND or HNC or equivalent”, “other professional qualifications e.g. nursing, teaching”, “none of the above” and “prefer not to answer”. We estimated years of education based on the highest qualification attained according to the International Standard Classification of Education mappings for United Kingdom^38^, and coded college or university degree as 16 years of education, A-levels as 13 years of education, O-levels/GCSEs as 11 years, CSEs as 11 years, NVQ/HND/HNC as 15 years, other professional training as 15 years, and the remaining two options coded as missing.

### Data processing and statistical analysis

We performed all data processing and analyses in Stata/SE 15.1 (StataCorp, College Station, TX, USA) on the Dementias Platform UK Data Portal^39^ using data from UK Biobank application 15008. Descriptive statistics were computed for selected sample characteristics, and potential sex differences were tested using the Mann-Whitney *U* test. Bivariate correlations were computed using Pearson’s correlation.

Structural equation models (SEMs) with maximum likelihood estimation were developed to test the patterns of association between ASI, CIMT, depression and cognition, and the potential mediating role of WMH. We performed this using a three-step procedure. First, the measurement models for the latent depression and cognition variables were tested. Next, the direct effects of ASI and CIMT on depression and cognition controlling for age and years of education were tested separately for men and women (Models 1a and b). Thereafter, we added WMH volume as a mediator between the vascular ageing variables and the outcome variables (Models 2a and b). The mediation model was considered significant if any of the associations attenuated with inclusion of the mediator. All continuous variables were mean-centred, and the scores on the verbal-numerical reasoning and reaction time tasks were multiplied by -1 for consistent directionality with higher values reflecting better cognition before entered into the SEMs.

### Patient and public involvement

Patients or members of the public were not involved in the design, recruitment, or conduct of this study, nor will they be involved in choosing the methods for dissemination of the results.

## Results

### Participants

This study included 7,394 participants aged 45 to 73 years, of which 54.8% are women. Table 1 presents the descriptive statistics for the sample. Tests for sex differences showed that overall, the women in this sample were significantly younger, had lower ASI, CIMT, and WMH burden, but performed worse on the reaction time and verbal numeric reasoning tasks and reported greater depression than men. Table 2 shows the bivariate correlations between the variables. ASI correlated with WMH and reaction, whereas CIMT correlated with WMH, reaction time, visual memory, and PHQ-9 scores. WMH correlated with all cognitive and depression measures.

**Table 1.** Descriptive statistics of sample characteristics (mean (SD)).

|  | Total (n=7,394) | Men (n=3,345) | Women (n=4,049) | *p* |
| --- | --- | --- | --- | --- |
| Age (years) | 61.13 (6.93) | 61.91 (7.01) | 60.49 (6.81) | <0.0001 |
| Arterial stiffness index | 9.57 (2.71) | 10.10 (2.62) | 9.13 (2.71) | <0.0001 |
| Carotid intima-media thickness (μm) | 663.65 (116.83) | 688.02 (130.51) | 643.51 (99.79) | <0.0001 |
| White matter hyperintensities volume (% ICV) | 0.29 (0.38) | 0.31 (0.39) | 0.28 (0.37) | <0.0001 |
| Intradividual mean reaction time (ms) | 578.91 (100.96) | 569.47 (98.72) | 586.71 (102.14) | <0.0001 |
| Visual memory score (incorrect matches) | 3.68 (2.92) | 3.71 (3.02) | 3.65 (2.84) | 0.7613 |
| Verbal-numeric reasoning score | 7.11 (2.05) | 7.32 (2.08) | 6.94 (2.01) | <0.001 |
| PHQ-9 total | 2.46 (3.33) | 2.08 (3.03) | 2.76 (3.53) | <0.0001 |

**Table 2.**
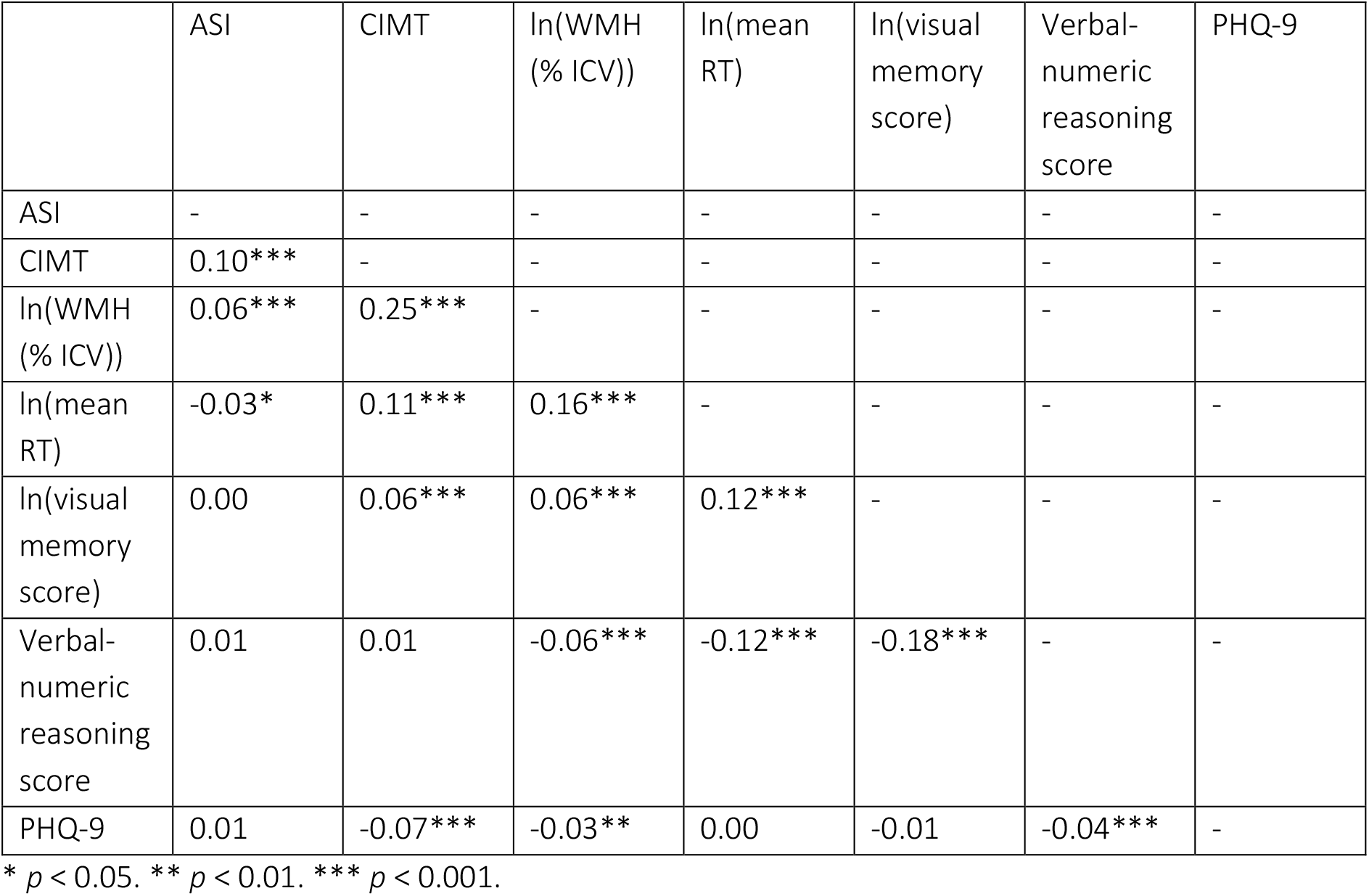
Bivariate correlations between the variables.

**Table 3.**
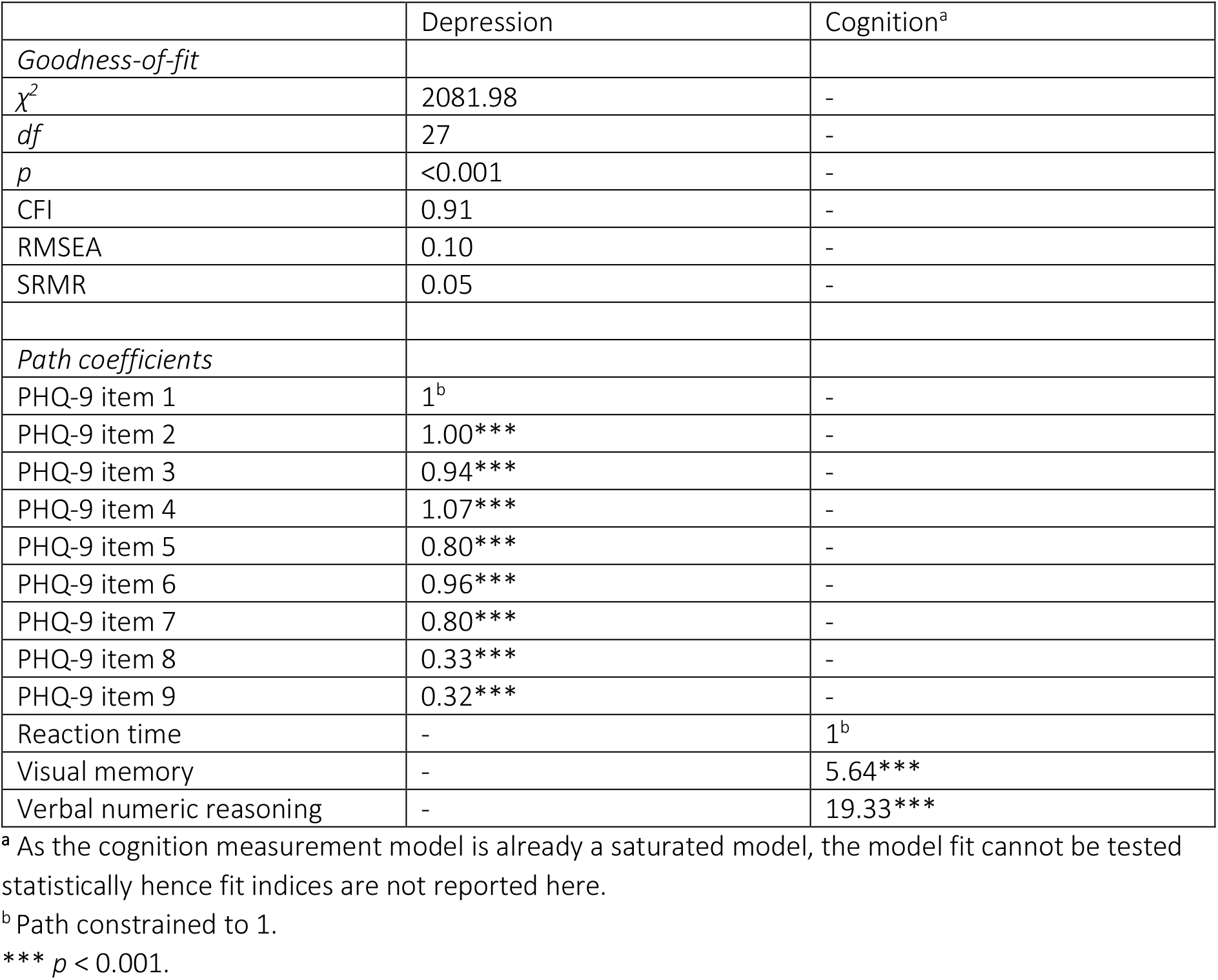
Fit indices and unstandardised path coefficients for the measurement models.

### Structural models

All four structural models showed good model fit (see Table 4). In men, there was a direct path between ASI and depression, and between depression and cognition (Model 1a). A higher ASI was associated with greater depression after controlling for age and years of education. Greater depression, in turn was associated with poorer cognition. The ASI-cognition indirect path, however, was non-significant. CIMT was not directly associated with either depression or cognition. Indirect paths linked CIMT to both depression and cognition in men (Model 2a). Greater CIMT was associated with higher WMH volume, which was associated with both greater depression and poorer cognition.

**Table 4.**
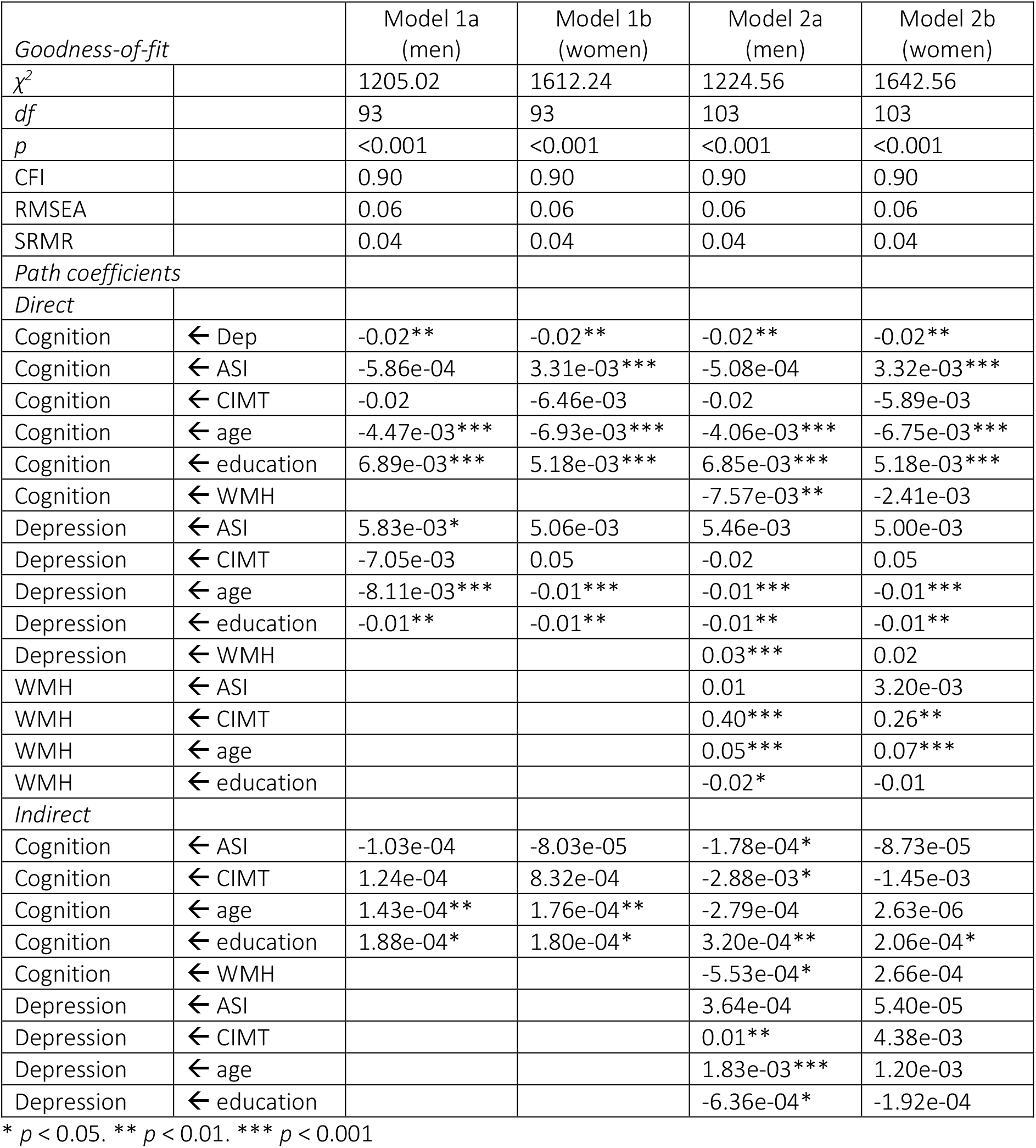
Fit indices and unstandardised path coefficients for the structural models.

| Goodness-of-fit |  | Model 1a<br>(men) | Model 1b<br>(women) | Model 2a<br>(men) | Model 2b<br>(women) |
| --- | --- | --- | --- | --- | --- |
| $\chi^2$ | | 1205.02 | 1612.24 | 1224.56 | 1642.56 |
| *df* |  | 93 | 93 | 103 | 103 |
| *p* |  | <0.001 | <0.001 | <0.001 | <0.001 |
| CFI |  | 0.90 | 0.90 | 0.90 | 0.90 |
| RMSEA |  | 0.06 | 0.06 | 0.06 | 0.06 |
| SRMR |  | 0.04 | 0.04 | 0.04 | 0.04 |
| *Path coefficients* |  |  |  |  |  |
| *Direct* |  |  |  |  |  |
| Cognition | ← Dep | -0.02** | -0.02** | -0.02** | -0.02** |
| Cognition | ← ASI | -5.86e-04 | 3.31e-03*** | -5.08e-04 | 3.32e-03*** |
| Cognition | ← CIMT | -0.02 | -6.46e-03 | -0.02 | -5.89e-03 |
| Cognition | ← age | -4.47e-03*** | -6.93e-03*** | -4.06e-03*** | -6.75e-03*** |
| Cognition | ← education | 6.89e-03*** | 5.18e-03*** | 6.85e-03*** | 5.18e-03*** |
| Cognition | ← WMH |  |  | -7.57e-03** | -2.41e-03 |
| Depression | ← ASI | 5.83e-03* | 5.06e-03 | 5.46e-03 | 5.00e-03 |
| Depression | ← CIMT | -7.05e-03 | 0.05 | -0.02 | 0.05 |
| Depression | ← age | -8.11e-03*** | -0.01*** | -0.01*** | -0.01*** |
| Depression | ← education | -0.01** | -0.01** | -0.01** | -0.01** |
| Depression | ← WMH |  |  | 0.03*** | 0.02 |
| WMH | ← ASI |  |  | 0.01 | 3.20e-03 |
| WMH | ← CIMT |  |  | 0.40*** | 0.26** |
| WMH | ← age |  |  | 0.05*** | 0.07*** |
| WMH | ← education |  |  | -0.02* | -0.01 |
| *Indirect* |  |  |  |  |  |
| Cognition | ← ASI | -1.03e-04 | -8.03e-05 | -1.78e-04* | -8.73e-05 |
| Cognition | ← CIMT | 1.24e-04 | 8.32e-04 | -2.88e-03* | -1.45e-03 |
| Cognition | ← age | 1.43e-04** | 1.76e-04** | -2.79e-04 | 2.63e-06 |
| Cognition | ← education | 1.88e-04* | 1.80e-04* | 3.20e-04** | 2.06e-04* |
| Cognition | ← WMH |  |  | -5.53e-04* | 2.66e-04 |
| Depression | ← ASI |  |  | 3.64e-04 | 5.40e-05 |
| Depression | ← CIMT |  |  | 0.01** | 4.38e-03 |
| Depression | ← age |  |  | 1.83e-03*** | 1.20e-03 |
| Depression | ← education |  |  | -6.36e-04* | -1.92e-04 |

\* $p < 0.05$ . \*\* $p < 0.01$ . \*\*\* $p < 0.001$

In women, there was a direct path between ASI and cognition, and between depression and cognition (Model 1b). A higher ASI was associated with better cognition after controlling for age and years of education. Similar to what was observed in men, greater depression as associated with poorer cognition. Again, CIMT was not directly associated with either depression or cognition. The mediation model showed that WMH did not mediate the ASI-cognition association in women (Model 2b).

## Discussion

Here we investigate potential sex differences in the associations between vascular ageing markers (ASI and CIMT), depression and cognition in 7,394 UK Biobank participants using structural equation modeling. The rationale for conducting the analyses stratified by sex is that previous research reported men and women show differences in the degree of arterial stiffness and CIMT in ageing. In this study, we found different cross-sectional patterns of association between men and women. ASI was directly related to different outcomes in men and women, with a higher ASI associated with greater depression in men, and rather intriguingly, with better cognition in women after controlling for the effects of age and education. We think the anomalous finding that higher ASI was associated with better cognition in women may be due to residual confounding, the source of which we were unable to identify. Contrary to earlier literature, we observed that CIMT was not directly related to depression or cognition in either sex. When WMH burden was taken into account, CIMT was related to WMH burden in both men and women, but WMH burden mediated the relationships between CIMT and depression and between CIMT and cognition only in men. ASI was not associated with WMH in either sex. Taken together, our findings add to the existing evidence base of sex differences in brain microvascular changes, depression and cognition in ageing, and suggest that they may be partly explained by sex-specific effects of vascular ageing.

Studies investigating arterial stiffness or CIMT in relation to WMHs, depression or cognition typically adjust for sex in their analyses^7–9 40–43^. The few studies that examined sex-specific associations reported mixed results. For example, some studies reported no association between CIMT and WMHs^44^ or between common CIMT and cognition in either of the sexes^45^. However, a small cross-sectional study of 91 women showed that increased CIMT was linked to poorer memory but not cognitive speed, and linked to both poorer memory and cognitive speed at 12-year follow-up^46^.

Interaction effects between sex and a range of risk factors may all play a role in these observed sex differences. For instance, hypertension, self-reported heart disease and high homocysteine levels were associated with WMHs in men^47^, whereas current smoking, lower high-density lipoprotein cholesterol and apolipoprotein A-1 levels were associated with WMHs in women^47 48^. Moreover, there is some evidence to suggest age x sex interactions may underlie differences in white matter ageing. For instance, different patterns of age-related white matter lesion localisation were observed between women and men^49^, and age-related reductions in regional white matter water diffusion metrics (i.e. cingulum fractional anisotropy, cingulum mean diffusivity and uncinate mean diffusivity) were also greater in men than in women in midlife^50^. Furthermore, the *APOE* ε4 allele, a known risk factor for sporadic Alzheimer’s disease^51^ as well as late-life depression^52^, also shows interaction effects with sex on a range of Alzheimer’s disease markers including beta-amyloid plaques^53^, neurofibrillary tangles^53 54^, cerebrospinal fluid levels of total tau and phosphorylated tau^55^, and hippocampal atrophy^56^. As these interaction effects may also result in different lesion localisation, they may explain the sex differences in mood and cognitive symptoms in ageing.

Given arterial stiffness and CIMT are considered early markers of vascular ageing, and may be mediators of the relationship between vascular risk factors and WMHs, depression or cognition, a better understanding of the mechanisms underlying the observed sex differences may have important prognostic or preventive implications.

Certainly, there are some limitations that need to be considered. The cross-sectional nature of this study precludes us from making any causal claims, it is possible that the observed sex differences relate to differences in the timing of when these changes occur. Further work is required to assess the casual role of vascular ageing changes in brain microvascular changes, depression and cognition as people age. The data for this study are from the imaging visit and the online follow-up, which began many years after the baseline assessment, and may show biased associations due to the “healthy volunteer effect”^57 58^. In addition, the majority of UK Biobank participants are Caucasian European and tend to be of higher socioeconomic status, which could have further limited the generalisability of the results.

Replication in an independent cohort would be beneficial, and longitudinal associations as well as white matter lesion localisation should also be examined in the future. If similar results were observed, the biological underpinnings of such sex-specific associations warrant further investigation, as a better understanding of these important sex differences could aid in the development of sex-specific preventive strategies for depression and cognitive decline or impairment in later life.

## Acknowledgements

This is a DPUK supported project with all analyses conducted on the DPUK Data Portal, constituting part 1 of DPUK Application 0132.

The Medical Research Council supports DPUK through grant MR/L023784/2.

## Contributions

All authors contributed to the conception and design of the study. RT analysed the data and drafted the manuscript. All authors were involved in the critical revision of the draft manuscript and approval of the final manuscript. The corresponding author attests that all listed authors meet authorship criteria and that no others meeting the criteria have been omitted.

## Transparency

The lead author affirms that the manuscript is an honest, accurate, and transparent account of the study being reported; that no important aspects of the study have been omitted; and that any discrepancies from the study as originally planned (and, if relevant, registered) have been explained.

