## Supplementary figures and images for "Sex differences in interrelationships between arterial stiffness, carotid intima-media thickness, white matter hyperintensities, depression and cognition: A UK Biobank study"

### Supplementary

Model 1a (men n=3,345):

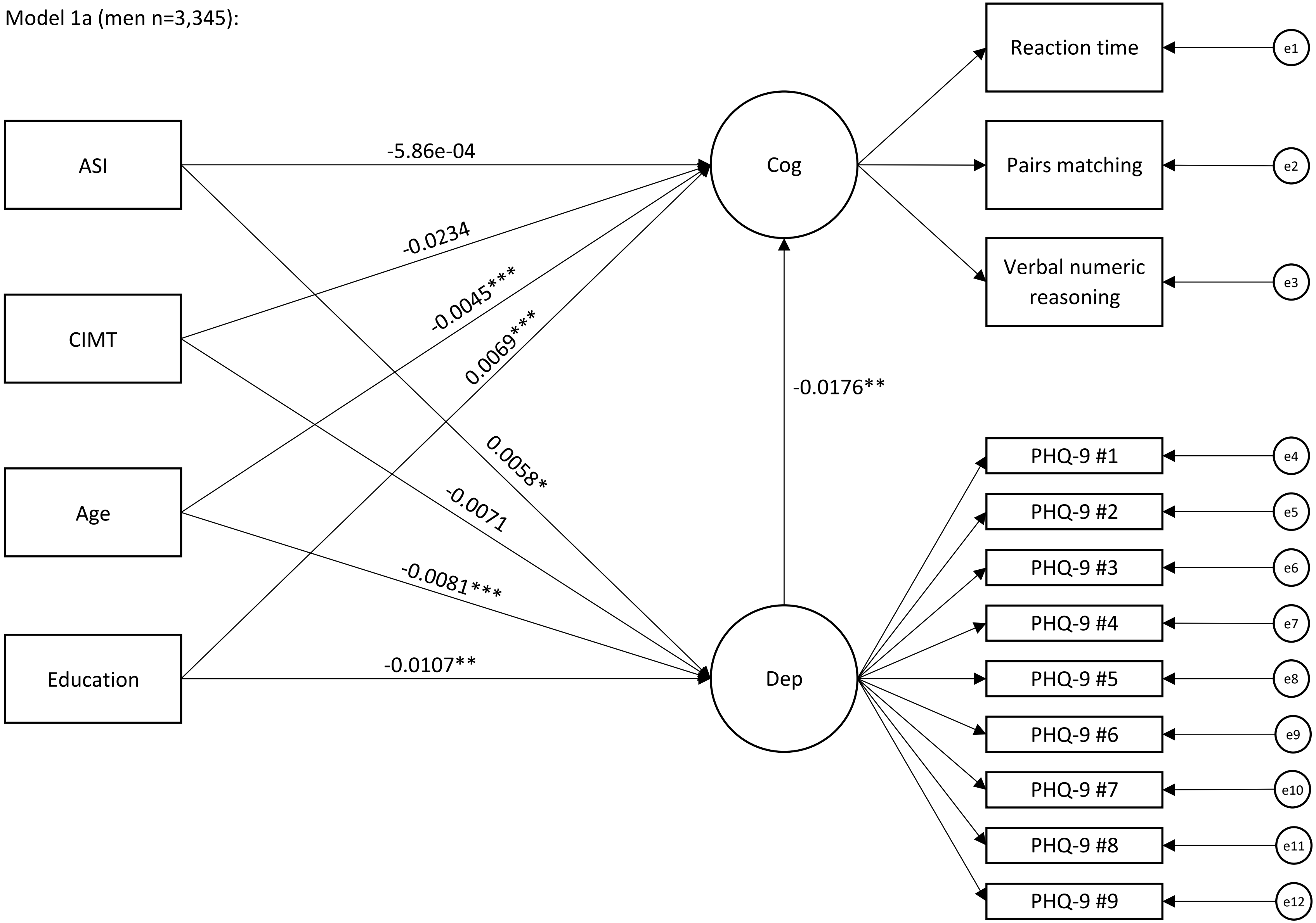

Model 1b (women n=4,049):

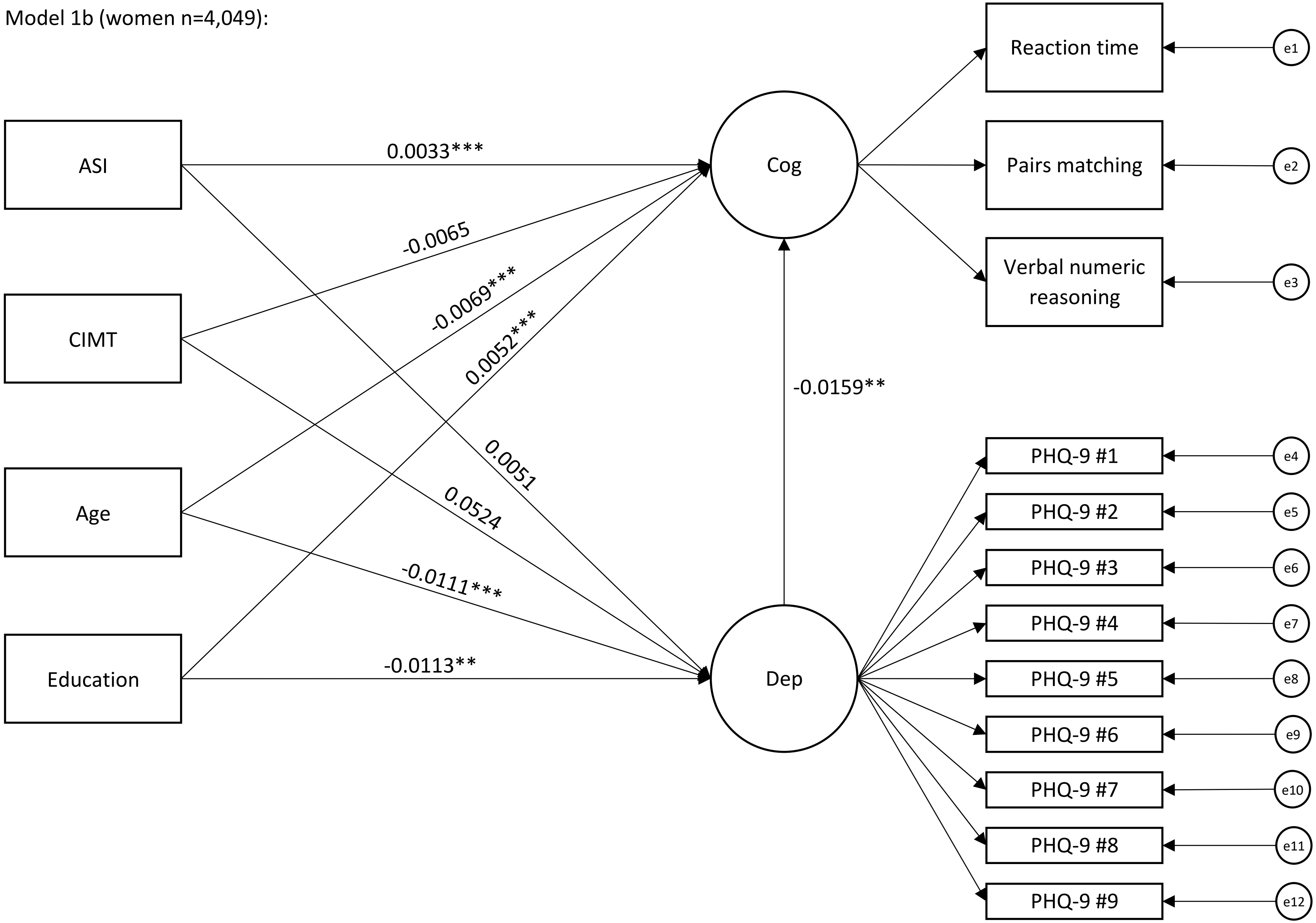

Model 2a (men n=3,345):

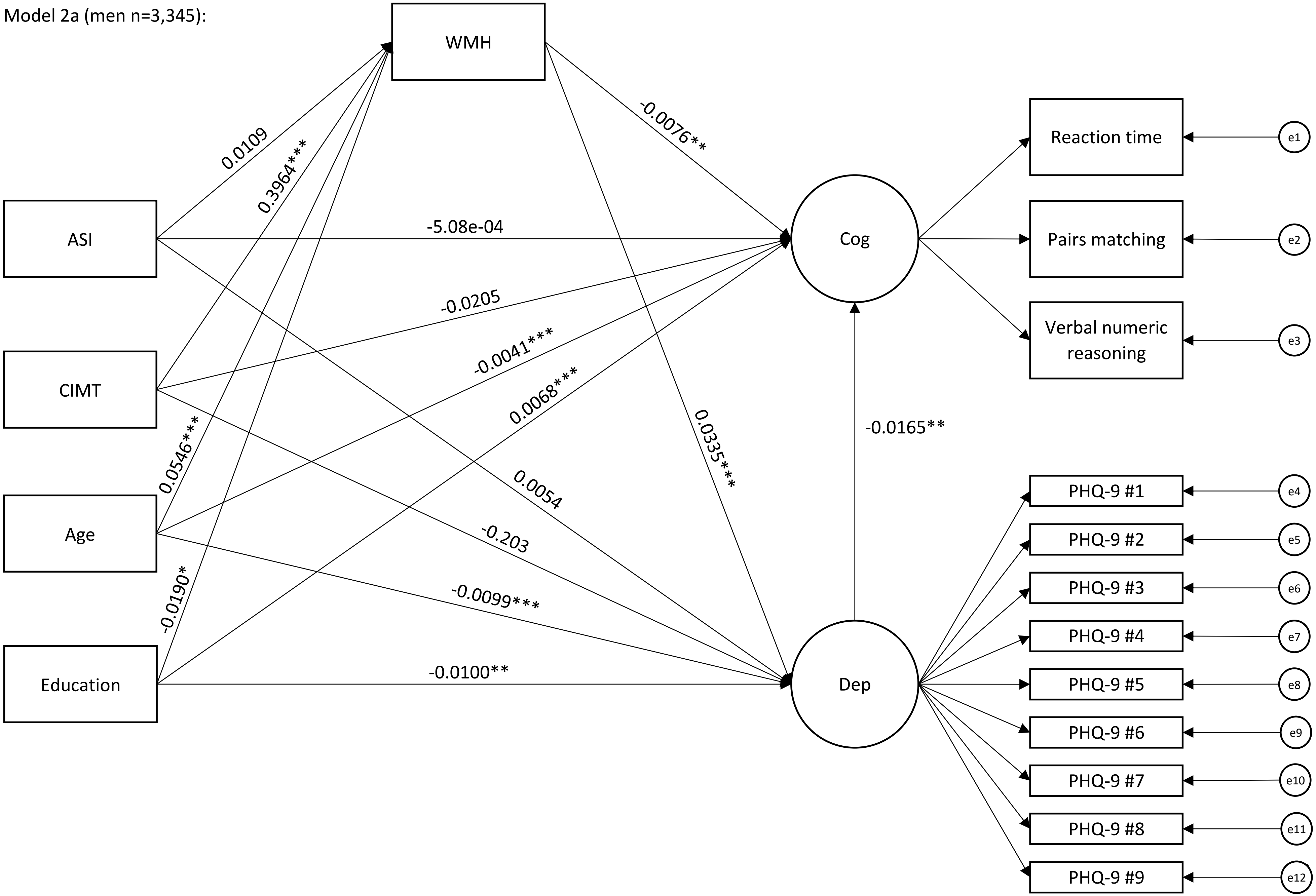

Model 2b (women n=4,049):

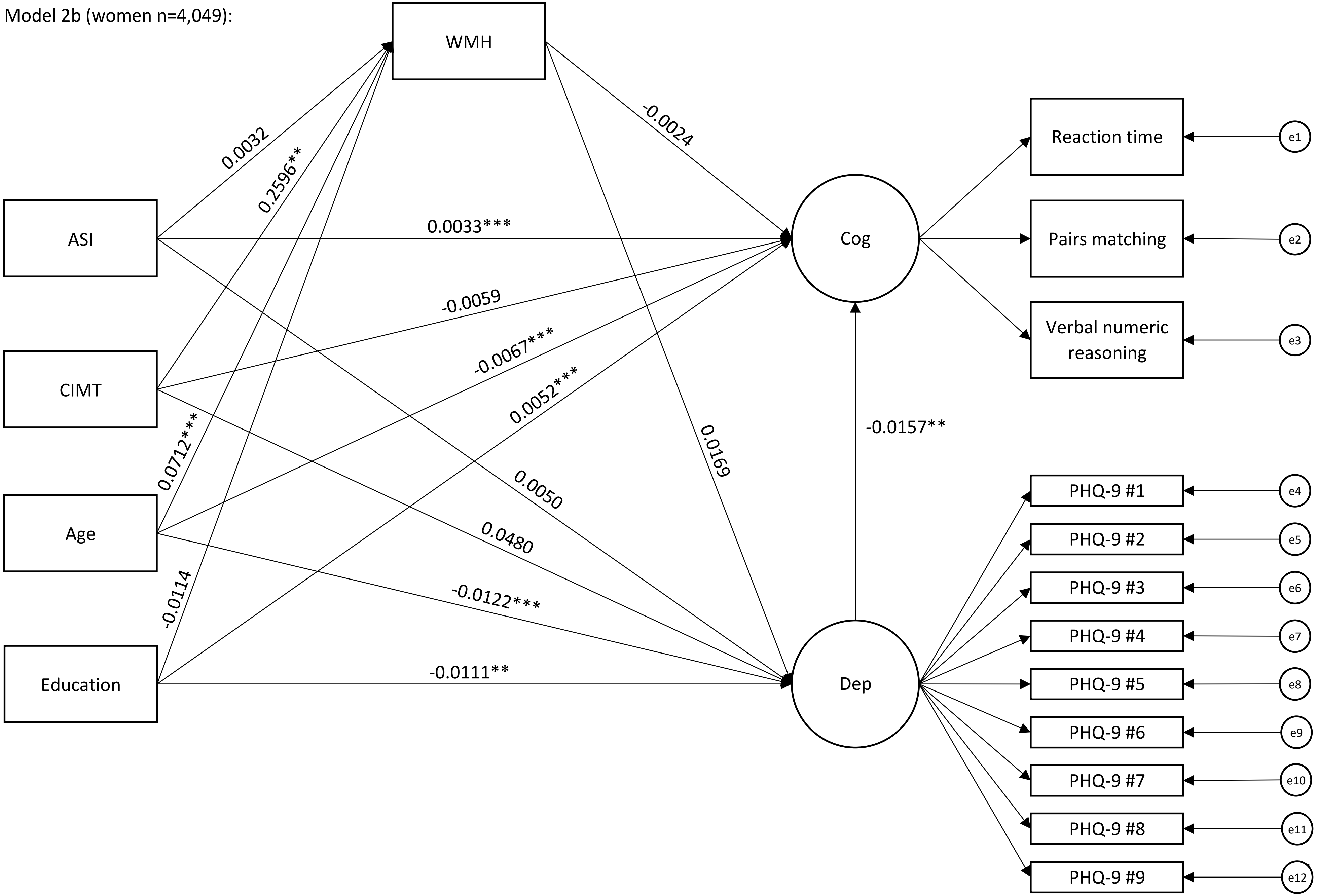
